# Intratumoral Microbiome of Adenoid Cystic Carcinomas and Comparison with other Head and Neck Cancers

**DOI:** 10.1101/2024.01.30.578054

**Authors:** Tatiana V. Karpinets, Yoshitsugu Mitani, Chia-Chi Chang, Xiaogang Wu, Xingzhi Song, Ivonne I Flores, Lauren K McDaniel, Yasmine M Hoballah, Fabiana J Veguilla, Renata Ferrarotto, Lauren E Colbert, Nadim J Ajami, Robert R Jenq, Jianhua Zhang, Andrew P Futreal, Adel K. El-Naggar

## Abstract

**Background:** Adenoid cystic carcinoma (ACC) is a rare, slow growing yet aggressive head and neck malignancy. Despite its clinical significance, our understanding of the cellular evolution and microenvironment in ACC remains limited.

**Methods:** We investigated the intratumoral microbiome of 50 ACC tumors and 33 adjacent normal tissues using 16S rRNA gene sequencing. This allowed us to characterize the bacterial communities within ACC and explore potential associations between the bacterial community structure, patient’s clinical characteristics, and tumor molecular features obtained through RNA sequencing.

**Results:** Bacterial composition in ACC displayed significant differences compared to adjacent normal salivary tissue and exhibited diverse levels of species richness. We identified two main microbial subtypes within ACC: oral-like and gut-like. Oral-like microbiomes, characterized by higher diversity and abundance of genera like *Neisseria, Leptotrichia, Actinomyces, Streptococcus, Rothia*, and *Veillonella* (commonly found in healthy oral cavities), were associated with the less aggressive ACC-II molecular subtype and improved patient outcomes. Notably, we identified the same oral genera in oral cancer and in head and neck squamous cell carcinomas. In both cancers, they were part of shared oral communities associated with more diverse microbiome, less aggressive tumor phenotype, and better survival. Conversely, gut-like microbiomes in ACC, featuring low diversity and colonization by gut mucus layer-degrading species like *Bacteroides, Akkermansia, Blautia, Bifidobacterium*, and *Enterococcus*, were associated with poorer outcomes. Elevated levels of *Bacteroides thetaiotaomicron* were independently associated with significantly worse survival, regardless of other clinical and molecular factors. Furthermore, this association positively correlated with tumor cell biosynthesis of glycan-based cell membrane components.

**Conclusions:** Our study uncovers specific intratumoral oral genera as potential pan-cancer biomarkers for favorable microbiomes in ACC and other head and neck cancers. These findings highlight the pivotal role of the intratumoral microbiome in influencing ACC prognosis and disease biology.

## INTRODUCTION

Adenoid cystic carcinoma (ACC) is a malignancy of major and minor salivary glands and less commonly of other organs [1]. It is slow growing aggressive tumor characterized by frequent local recurrence and delayed distant metastasis [2]. ACC is subdivided into 3 histological groups (tubular, cribriform, and solid) [3] and into 2 molecular subtypes, ACC-I and ACC-II. The former is characterized by activation of MYC and MYC target genes, enrichment of NOTCH-activating mutations, and significantly worse overall patient survival when compared with ACC-II. [4]. There is little data on factors underlying development and progression of the disease [2, 5]. A recent small study of saliva from 13 ACC patients and 10 healthy controls reported significant difference in taxonomic structure of salivary microbiome between these 2 groups with greater abundances of *Rothia* and *Streptococcus* in the saliva of ACC patients [6]. The findings highlight difference of the intratumoral microbiome from the normal tissue and suggest a role for salivary microbiome as a potential biomarker of ACC.

Although intratumoral microbiome has not been studied in ACC, numerous studies of bacterial communities in solid cancers including colorectal [7], oral [8], pancreatic [9, 10], and gynecological cancer [11] revealed difference in bacterial diversity and taxonomic composition between tumor and normal tissue. While challenges remain in definitively determining the abundance, roles, and consequences of intracellular bacteria in cancer [12], emerging evidence strongly suggests their potential for diagnosis, prognosis, and treatment of various tumors [13-15]. Both, positive and negative associations of intratumoral bacteria with tumor development, progression, patient survival, and response to therapies have been reported with different underlying molecular mechanisms [16] [17]. The studies linked certain microorganisms to prognosis and treatment of cancer including pancreatic [18], colorectal [7], and others [19].

An important function of salivary gland is the secretion of saliva that contains antimicrobial factors impacting microorganisms [20, 21] that may play a role in transformation of salivary epithelia and other cells in course of ACC development, progression, and treatrment. Salivary gland-derived mucins are important components mediating interactions of microorganisms with the epithelial cells and their environment [22]. Mucins are glycoproteins produced by epithelial cells and comprised of a protein and a glycan moiety that has recognition motifs and binding sites for microbes [23]. The interaction of microbes with mucins plays a central role in maintaining the oral health. It is not known, however, how the ACC development will impact microbial diversity, taxonomic organization, and interaction between microbes and salivary gland components.

In this study, we analyzed taxonomic structure and diversity of intratumoral microbiomes in ACC tumor and adjacent non-tumoral tissue using 16S RNA gene sequencing, explore associations of the microbiomes with clinical characteristics and molecular features inferred by RNA-sequencing, and compared intratumoral microbiomes in ACC with oral and head and neck cancers.

## METHODS

### Study design

The study was conducted in compliance with the Declaration of Helsinki and approved by the institutional review board. The study materials available for 16S rRNA gene sequencing was comprised of 50 fresh frozen ACC tumors (Fig. S1). Histologically non-tumoral adjacent tissue was available only for 33 of these patients. Eighty-eight samples were sequenced, which include 33 pairs of ACC tumor and adjacent non-tumoral tissue (66 samples), 17 additional tumor samples (without paired normal samples) and 5 additional samples that were duplicated to evaluate potential batch effect (Fig. S2). IDs of the sequenced samples available for download as raw sequencing reads from NCBI are provided in supplementary table (Table S1) with other clinical information of the patients.

### DNA extraction and 16S rRNA gene sequencing

Snap-frozen human tumor and normal specimens were processed and analyzed at the MD Anderson Cancer Center Microbiome Core Facility. Microbial genomic DNA was extracted using the DNeasy Powersoil Pro DNA kit (Cat No. 47014, QIAGEN), following the manufacturer’s instructions. The extraction process involved adding 100 mg to 200 mg of homogenized adenoid cystic carcinoma (ACC) or corresponding normal tissue to a bead beating tube for efficient homogenization and lysis using mechanical and chemical methods.

Subsequently, the lysed cells were treated with the Inhibitor Removal Technology (IRT) solution to remove inhibitors. The total genomic DNA was then captured on a silica membrane in a spin-column format, followed by washing and elution steps. The Earth Microbiome Project [24] method was adapted to generate amplicon libraries of the V4 hypervariable region of the 16S rRNA gene. PCR amplification of microbial DNA was carried out using 515F and 806R primer constructs containing sequencing-ready barcodes and adapter sequences. The quality and quantity of the barcoded amplicons were assessed using an Agilent 4200 TapeStation system (Agilent) and Qubit Fluorometer (Thermo Fisher Scientific). The amplicons were pooled in equimolar ratios. The pooled libraries were quantified using a Qubit fluorometer, and their molarity was calculated based on the amplicon size. Sequencing was performed on the Illumina MiSeq platform (Illumina) using the 2x250 bp paired-end protocol, resulting in paired-end reads with near-complete overlap. The custom sequencing primers used were as follows: Read1 sequencing primer: 5’-TATGGTAATTGTGTGYCAGCMGCCGCGGTAA-3’; Read2 sequencing primer: 5’-AGTCAGCCAGCCGGACTACNVGGGTWTCTAAT-3’; and index sequencing primer: AATGATACGGCGACCACCGAGATCTACACGCT. The sequencing data from the paired-end reads were de-multiplexed using QIIME [25]. The paired-end reads were merged, followed by dereplication and length filtering using VSEARCH v7 [26]. De-noising and chimera calling were performed using the unoise3 command [27]. Bacterial taxonomies were assigned using the SILVA database version 138 (https://www.arb-silva.de/) [28].

### Statistical analyses

Bacterial diversity was calculated at the level of predicted OTUs and at the genus level in terms of different diversity measures using R library ‘microbiome’ [29]. Differentially abundant taxa between 33 paired tumor and adjacent normal tissue were determined by 2 different tools, MaAsLin2 [30] and by LeFSe (Logarithmic Discriminant Analysis Effect Size) [31] using default parameters. In case of LeFSe, the logarithmic discriminant analysis (LDA) score threshold was set to 1.5 to identify more differentially expressed taxa.

Supervised hierarchical clustering of most common OTUs identified in ACCs was implemented by open source clustering software Cluster 3 [32] with default parameters using centroid linkage as the clustering method. Samples were ordered by abundance of the *B. thetaiotaomicron* and by sum of 3 species abundances, *Granulicatella adiacens*, unclassified *Leptotrichia*, and *Rothia mucilaginosa* for *B thetaiotaomicron* negative samples.

Permutational multivariate analysis of variance (PERMANOVA) and nonparametric multivariate analysis of variance (MANOVA) [33] were applied to 33 normal and tumor pairs to test if the groups have distinct taxonomic composition and to 50 ACC tumor samples to test association of the proposed grouping of the samples with taxonomic composition. The analysis was implemented using R libraries “vegan” [34] (https://rdrr.io/cran/vegan/, R package version 2.6-4), functions adonis2() and permutest(). Defaul parameters were used if it is not specified differently in the text.

Associations with survival was explored for most common and abundant OTUs identified in 50 ACC tumors using the Cox proportional-hazards model: coxph() function in R library “survival” [35] and R library “survminer” [36]. Namely, we aggregated the produced OTU table at the species level and selected those OTUs that are most common in ACC tumors (found in more than 20 samples out of 50). Significance of associations was evaluated by log-rank, likelihood test, and Wald test p-values. Bonferroni correction [37] was used to adjust the p-values.

Gene set enrichment analysis (GSEA) was implemented for 44 samples with available RNA-seq data using R package “fgsea” (fast pre-ranked GSEA) [38] downloaded from https://github.com/ctlab/fgsea. Ranking of genes for the analysis was done using Spearman correlation calculated between *B theta* abundances and the gene expression for each gene across 44 samples. The gene sets representing KEGG pathways (https://www.gsea-msigdb.org/gsea/msigdb/human/genesets.jsp?collection=CP:KEGG) [39] and Gene Ontology Biological processes (https://www.gsea-msigdb.org/gsea/msigdb/human/genesets.jsp?collection=GO:BP) [40] were downloaded from the Molecular Signature Database v2022.1 [41].

Comparison of ACC bacterial community structure with the oral cancer was implemented using a recently published cohort of 33 oral cancer patients with tumors analyzed by 16S RNA gene sequencing [8]. The OTU table was aggregated to the *Genus* level in each of the 2 cohorts, referred as Oral and ACC, and *Genera* that are most common (found in 75% of samples) were selected for unsupervised hierarchical clustering implemented separately for each cohort by open source clustering software Cluster 3 [32] with default parameters using centroid linkage as the clustering method.

Comparison of ACC with the head and neck squamous cell carcinoma was implemented using predicted bacterial genera in TCGA-HNSC cohort. The dataset is described by Gihawi et. Al [42] and available for download as supplementary table. Briefly, the OTU table was created by re-filtering the unmapped reads _b_y aligning them to the human CHM13 reference genome and by matching the re-filtered reads with a curated Kraken database [43]. We selected only primary tumors, fresh frozen, analyzed by WGS using Illumina HiSeq for our comparison. The OTU table of the obtained 170 samples was further filtered by including 50 topmost common bacterial species to reduce scarcity of the table for unsupervised hierarchical clustering, which was implemented as described above. Clinical information of the patients was downloaded from TCGA using R library ‘TCGAbiolinks’ [44].

## RESULTS

### Clinical characteristics of the cohorts

Fresh frozen primary tumors from 50 patients with ACC, stage 1-4, with no prior therapy at time of surgery, were used for 16S RNA gene sequencing to characterize intratumoral bacterial composition, diversity, structure, and association with the patient clinicopathological factors (Fig. S1). Thirty-three patients had also adjacent non-tumoral tissue sequenced for comparative analysis of microbial communities with those in the tumor. A subset of 44 patients (88%) had their tumors analyzed using RNA-seq [4], and a classification of these tumors into 2 molecular subtypes, referred as ACC-I and ACC-II, was available for correlative analysis with the intratumoral microbiome. Patients with both molecular subtypes (45% ACC-I and 55% ACC-II) were present in the study cohort (Table 1, Supplementary table). Primary sites of ACC in the cohort were in maxillary sinus, base tongue, palate, parotid, trachea, sublingual, lacrimal gland, and submandibular. The sites are close to oral, throat, nasal, and eye ocular surface body sites, characterized by distinct microbiomes according to Human Microbiome Project. Most of the patients were males (66%), had stage 3-4 tumors at diagnosis (70%), solid or cribriform histology (84%), and perineural invasion (PNI) (84%). All the patients underwent primary tumor resection, and most of them were treated with postoperative RT (88%) and/or chemotherapy (38%).

**Table 1.**
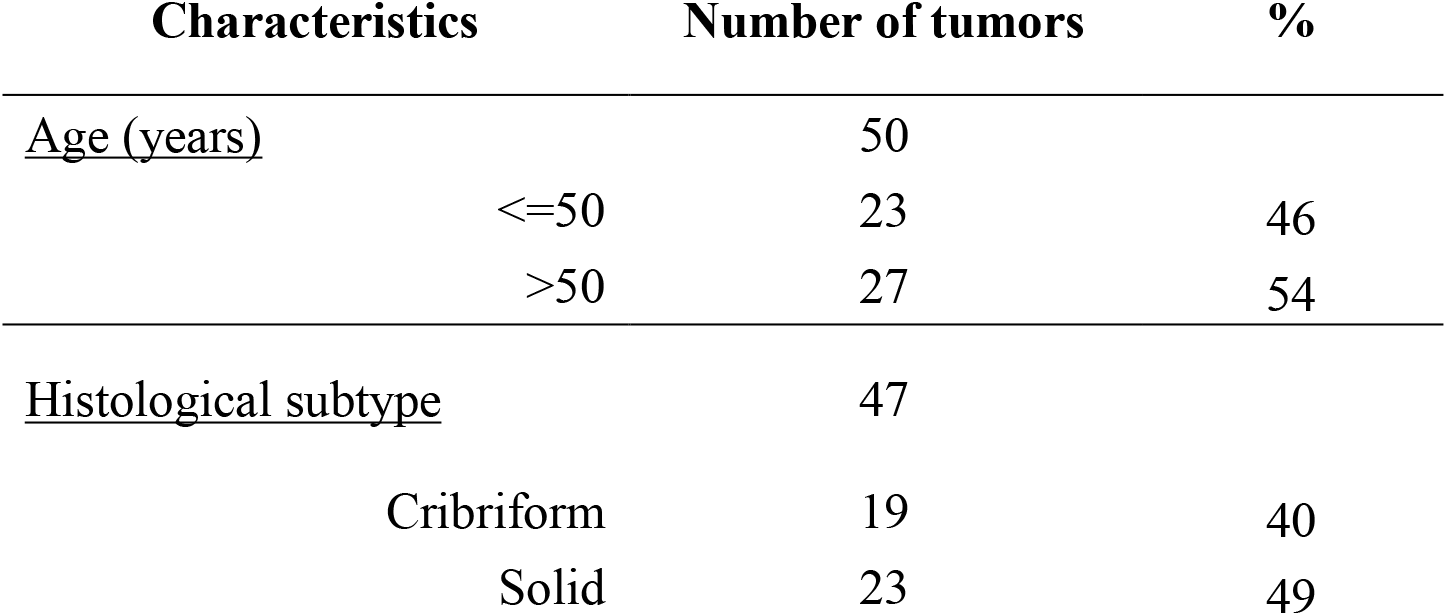
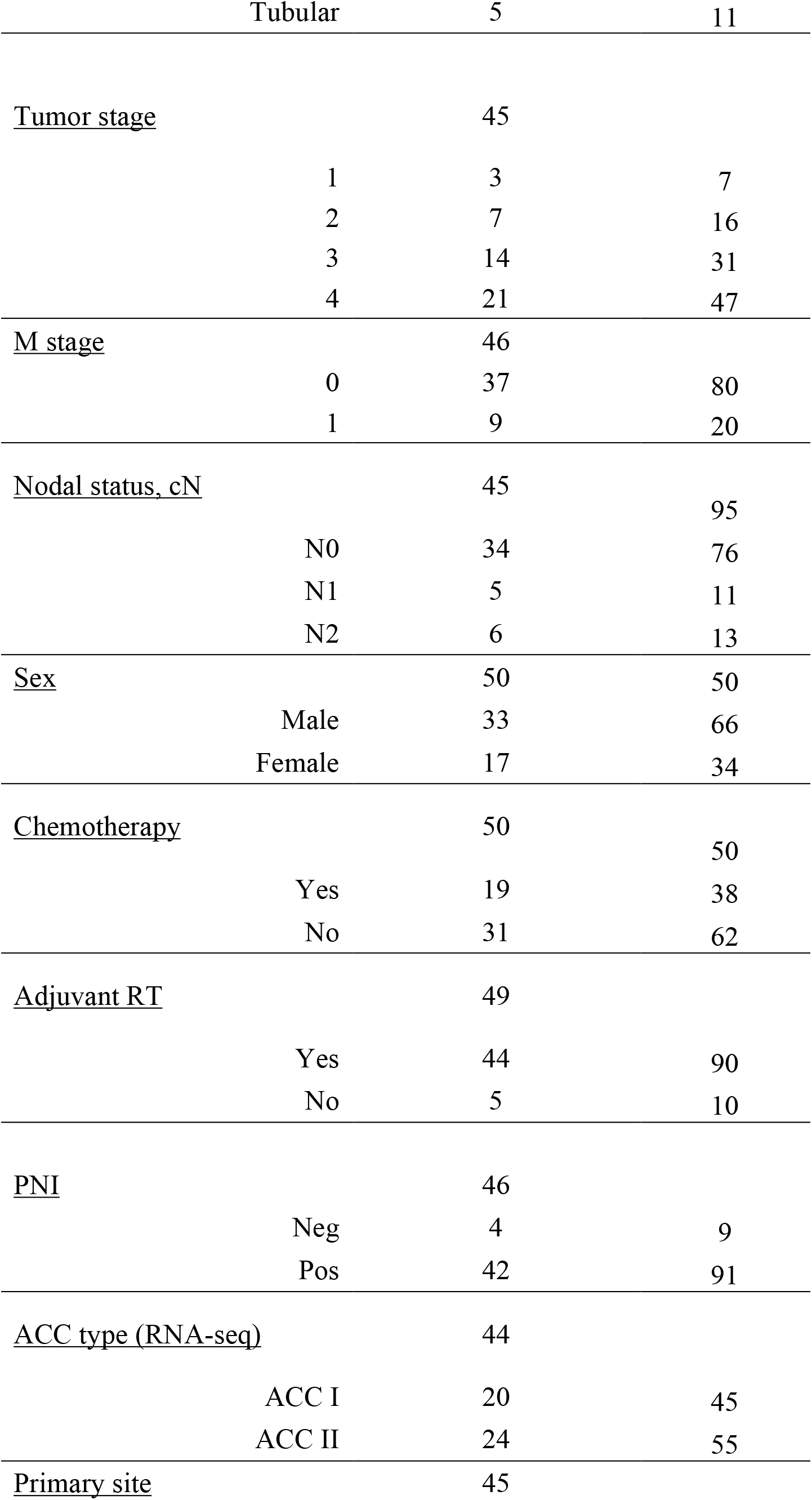
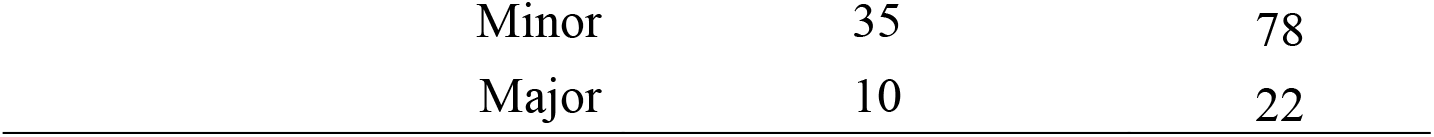
Clinicopathological characteristics of the ACC cohort

| Characteristics | Number of tumors | % |
| --- | --- | --- |
| <u>Age (years)</u> | 50 |  |
| <=50 | 23 | 46 |
| >50 | 27 | 54 |
| <u>Histological subtype</u> | 47 |  |
| Cribriform | 19 | 40 |
| Solid | 23 | 49 |

|  |  |  |  |
| --- | --- | --- | --- |
|  | Tubular | 5 | 11 |
| <u>Tumor stage</u> |  | 45 |  |
|  | 1 | 3 | 7 |
|  | 2 | 7 | 16 |
|  | 3 | 14 | 31 |
|  | 4 | 21 | 47 |
| <u>M stage</u> |  | 46 |  |
|  | 0 | 37 | 80 |
|  | 1 | 9 | 20 |
| <u>Nodal status, cN</u> |  | 45 |  |
|  |  |  | 95 |
|  | N0 | 34 | 76 |
|  | N1 | 5 | 11 |
|  | N2 | 6 | 13 |
| <u>Sex</u> |  | 50 | 50 |
|  | Male | 33 | 66 |
|  | Female | 17 | 34 |
| <u>Chemotherapy</u> |  | 50 |  |
|  |  |  | 50 |
|  | Yes | 19 | 38 |
|  | No | 31 | 62 |
| <u>Adjuvant RT</u> |  | 49 |  |
|  | Yes | 44 | 90 |
|  | No | 5 | 10 |
| <u>PNI</u> |  | 46 |  |
|  | Neg | 4 | 9 |
|  | Pos | 42 | 91 |
| <u>ACC type (RNA-seq)</u> |  | 44 |  |
|  | ACC I | 20 | 45 |
|  | ACC II | 24 | 55 |
| <u>Primary site</u> |  | 45 |  |
| Minor | 35 | 78 |
| Major | 10 | 22 |

### Taxonomic structure of the bacterial community in ACC tumors and in adjacent normal tissue is associated with bacterial richness

The 16S RNA gene sequencing of 50 ACC tumors identified median number of 200 putative bacterial species per sample, ranging from 52 to 673 OTUs. Further paired comparison of 33 tumors and normal adjacent tissue in terms of diversity metrics revealed significantly increased species richness (not adjusted two-tailed paired t-test P value is 0.005) and related characteristics, Fisher’s diversity and Chao1 index (P=0.005), in normal samples. No significant differences (P>0.05) between normal and tumor tissue were found in other indexes of diversity. Further analysis showed significant pairwise correlation (R=0.73, P<1E-5) of the species richness between tumors and paired normal tissues (Fig. S2). Significant decrease in the number of species within the tumor was observed only in patients with rich microbiome; no association was found when tissue had low richness (Fig 1A, Fig. S3). There was also a trend (Fisher test P=0.09) for females rather than males to have richer microbiome in both, tumor and normal tissue.

**Fig. 1.**
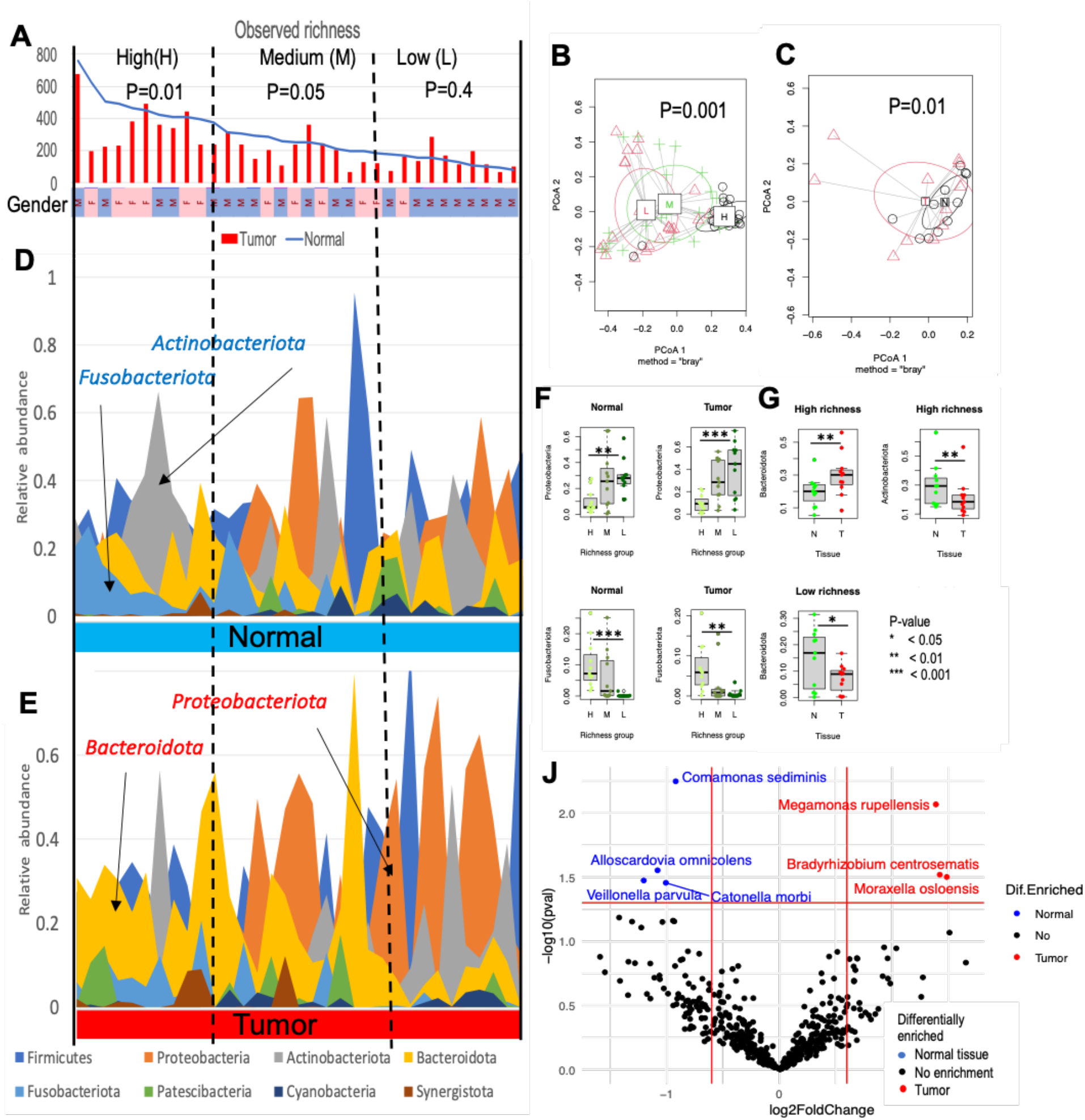
Associated changes in richness and taxonomic structure of bacterial communities in tumor and matched normal tissue. **A**. Significant difference in number of identified spp (richness) in tumors vs normal tissue is mostly observed in patients with more diverse microbiome in normal tissue. Samples were sorted by richness and then divided into 3 equal richness groups (see dashed lines): with low (L), medium (M), and high (H) number of species in the normal samples. Paired two-sided t-test was used to evaluate if richness is significantly different between tumor and normal tissue in each group. The produced p-values from the test are shown on the figure. Gender of the patient is labelled as a bar at Y axis by rose (female) and blue (male) color. **B**. Permutational multivariate analysis of variance of paired tumor-normal tissue in terms of bacterial species abundances reveals significant association of the bacterial community structure with the microbiome richness. Centroids of samples for each richness group are labelled as H (High richness), M(Medium), and L(Low). **C**. Permutational multivariate analysis of variance of paired tumor-normal samples in patients from the High richness group reveal significant difference in the community structure between tumor and normal tissue. Centroids for tumors (red triangles) and normal (black circles) samples are labelled by T and N respectively. **D. E**. Relative abundance of top abundant phyla in normal and tumor samples sorted by species richness in normal tissue. Phyla that are significantly more abundant in normal or in tumor tissues are labelled by blue and red arrows accordingly. Increased abundance of *Proteobacteriota* and decreased abundance of *Actinobacteriota* and of *Fusobacteriota* are observed in tumors with low species richness. **F**. Significant difference in *Bacteroidota* and *Actinobacteriota* relative abundances between tumor and normal tissue in patients from different richness group. While in tumors with rich microbiomes, *Bacteroidota* relative abundances is increased, in low richness microbiomes, the relative abundance of the phylum is decreased. Only tumors from high richness group have significantly decreased *Actinobacteriota* abundances. **J**. Volcano plot of differentially abundant species identified by MaAsLin in normal and tumor tissue.

The taxonomic structure of the bacterial community in tumor and normal samples also depended on species richness. Applying permutational multivariate analysis of variance to all 33 paired tumor-normal samples, we observed significant difference in taxonomic composition when the paired samples was divided into 3 equal groups according to normal tissue richness: high (H), medium (M), and low (L) (Fig. 1B). The community structure of the H group samples differed from the M group and, especially, the L group at both the OTU level (Fig. 1B) and the *Phylum* level (Fig. S4B). No difference was found between tumor and normal tissues when analyzing all samples together (Fig. S4A). However, when examining each group separately, a significant difference in taxonomic structure between tumor and normal tissue was observed in the H richness group of samples (Fig. 1C and Fig. S4C), but not in the M or L group (Fig. S4D). Due to inconsistent changes between the groups, more significant differences in relative abundances of *Phyla* were observed if samples in L and H richness groups were considered separately (Fig. 1F, Fig. 1G, Fig. S5, and Fig. S6). Normal and tumor tissue with low versus high species richness showed significant increase in the abundance of *Proteobacteriota* (P=0.001 and P=0.0007, Fig. 1F) and significant decrease in *Fusobacteriota* (P=0.0003 and P=0.002, Fig. 1F). Tumors versus normal in H group (with high number of OTUs) showed increase of oral *Bacteroidota* and decrease of oral *Actinobacteriota* (Fig. 1G), and tumors versus normal in L group (low number of OTUs) showed decreased abundance of *Bacteroidota* (Fig. 1G) and increased abundance of *Proteobacteriota* (Fig. S6).

Association of the taxonomic structure with richness was also confirmed at the *Order* level (Fig. S7) and the OTU level (Fig. S8, Fig. 1J, Fig. S9). Differentially abundant OTUs between normal and tumor tissue were found by comparison of 33 paired ACC tumor-normal tissues using MaAsLin2 (Fig. 1J) and LeFSe (Fig S8, Fig S9, Fig. S10). Four putative species, including typical inhabitants of human oral cavity *Veillonella parvula [45], Catonella morbi [46]* and *Alloscardovia omnicolens* [47] were abundant in normal tissue (Fig 1J) and, specifically, in samples with total number of species above the median value (Fig. S9). Opposite relationship with species richness was found for 2 putative species differentially abundant in tumors (Fig S10): a gut bacterium *Megamonas rupellensis* [48] and *Bradyrhizobium centrosematis*, a known contaminant of purified and municipal water system [49]. Both species were abundant in tumor tissue with low species richness. *Moraxella osloensis* implicated in inflammation of ocular membranes [50] was also more abundant in tumor tissue but was not associated with the species richness.

### Association of bacterial composition and abundances with overall patient survival

To find clinical factors and specific intratumoral species that might be associated with survival outcome, we analyzed tumor tissue in 50 patients (Fig. S1). Analysis of clinical factors using the Cox proportional-hazards model identified only one confounder, the ACC molecular subtype (ACC-I versus ACC-II), significantly associated with survival after multiple testing correction (log-rank Padj = 0.003) (Table S2). Among diversity characteristics, only diversity inverse Simpson, diversity coverage, and Gini inequality index showed statistical significance (log-rank Padj<0.05), although they were not independent predictors of survival probability from the ACC molecular subtype (Table S2).

Of topmost common and abundant species analyzed by the Cox proportional-hazards model, only 1 OTU significantly (p-value is 8E-05) negatively associated with survival after adjustment of log-rank p-value for multiple testing (Fig 2A). The OTU was classified taxonomically with 100% identity as a well-known commensal resident of the human gut [51] *Bacteroides thetaiotaomicron (B. theta)*. Importantly, the abundance of *B theta* predicted survival probability independently from ACC subtype, and combination of *B. theta* abundance and ACC subtype increased significance of the model from P=2E-4 to P=3E-5 according to the likelihood ratio test (Table S3). Only 3 OTUs (Fig 2A, Table S2), taxonomically classified as typical inhabitants of the oral cavity *Granulicatella adiacens, Rothia mucilaginosa*, and an unclassified species of *Leptotrichia*, had significant positive association with survival (likelihood ratio test P, not adjusted, is 0.03, 0.05, and 0.05, respectively). Abundances of the OTUs significantly positively correlated with each other across the samples (rho is from 0.36 to 0.70) suggesting that the oral species can be a part of the same community. Each of the species had significant negative correlation with *B. thetaiotaomicron* abundance (rho is from -0.27 to -0.49) revealing their potential negative impact on outgrowth of *B. thetaiotaomicron*.

**Fig. 2.**
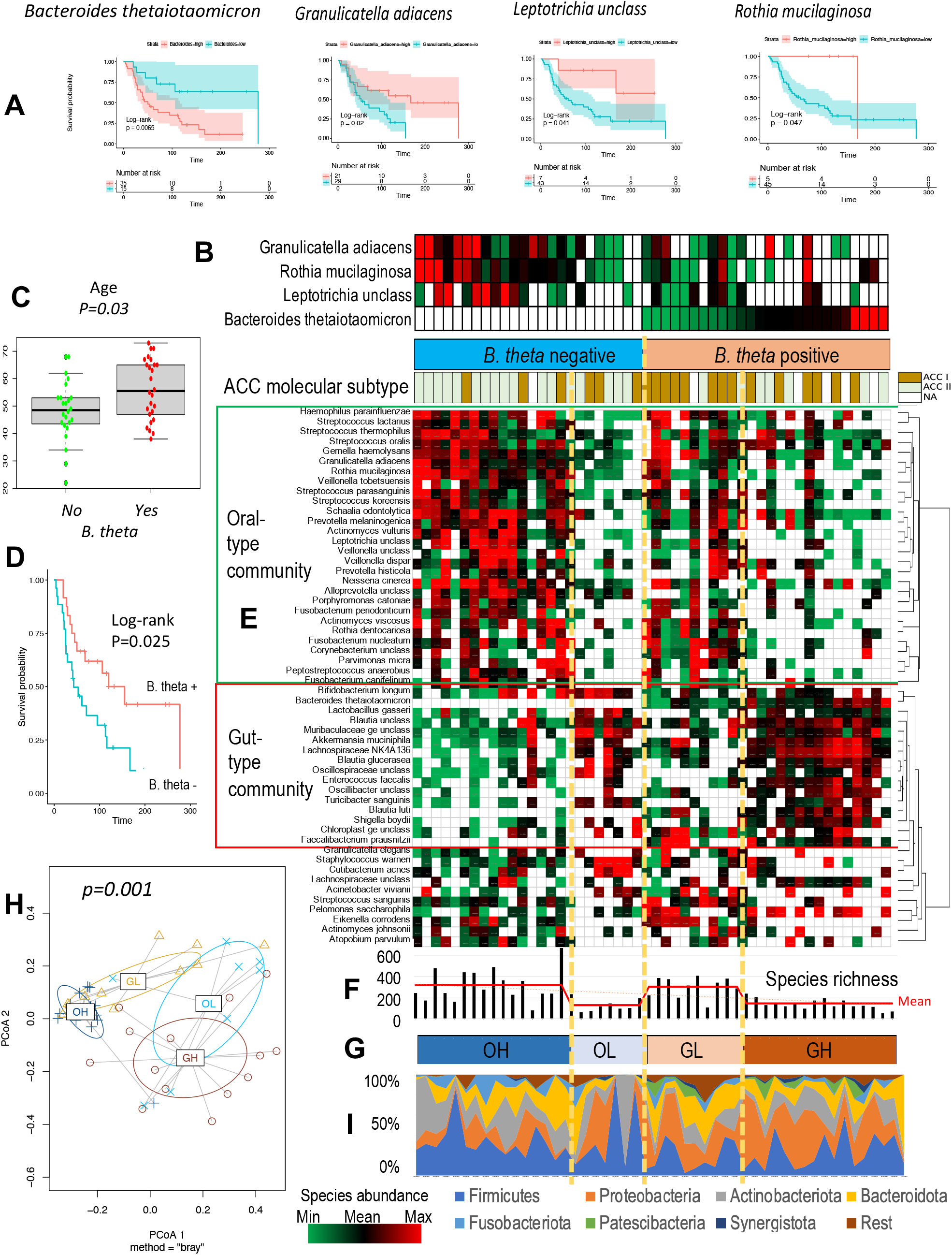
Grouping of ACC intratumoral microbiomes according to bacterial community type and abundance. **A**.Bacterial species significantly associated with overall survival. **B**.Grouping and ordering of ACCs by abundances of species associated with survival. The studied ACCs were subdivided into 2 groups, *B. theta* positive and negative. The former group was ordered by sum of 3 OS-positive spp abundance, from max to min and the latter by *B. theta* abundance from min to max. The *B. theta* positive group of ACC tumors were dominated by molecular subtype ACC-II, while ACC-I subtype was significantly more common in *B. theta* negative ACCs. **C**.Association of *B. theta-* and *B. theta+* tumors with age of ACC patient. **D**.Association of *B. theta-* and *B. theta+* tumors with ACC patient survival. **E**.Supervised hierarchical clustering of 54 most common spp identified in ACCs reveals Oral- and Gut-type bacterial communities associated with *B. theta* negative and positive tumor groups accordingly. Oral-type bacterial community is enriched with common oral species, while Gut-type community is enriched with species that are common for the human gut. **F, G**. Grouping of ACC intratumoral microbiomes according to Oral- and Gut-community abundances and association of the groups with richness; OH: High abundance of Oral spp, OL: Low abundance of Oral spp, GH: High abundance of Gut spp, GL: Low abundance of Gut spp. OH and GL microbiomes have significantly increased richness than OL and GL microbiomes. **H**. Permutational multivariate analysis of variance confirms significant difference in the microbial composition of proposed intratumoral microbiome groups (OH, OL, GH, and GL). **I**. Significant association of bacterial community taxonomic structure at the phylum level with proposed grouping of intratumoral microbiome in ACC. *Firmicutes, Actinobacteriota*, and *Fusobacteriota* are more abundant in *B. theta-* negative group, while phyla of *Proteobacteriota, Patescibacteria*, and *Cyanobacteria* are outgrowing in *B. theta+* tumors.

To explore associations of the species with other topmost common colonizers of intratumoral microbiome, we group tumors and species in terms of *B. theta* and the oral species abundances (Fig. 2B and 2C). We split tumors into *B. theta+* (positive) and *B. theta-* (negative) and then ordered *B. theta*+ tumors by *B. theta* abundance from max to 0 (from right to left) and *B. theta* negative samples by sum of the 3 oral species abundances from min to max (Fig. 2B). Patients with *B. theta-* tumors were younger (Fig. 2C), had better outcome (Fig. 2D), and their tumors more likely had ACC-II myoepithelial molecular subtype (Fisher’s test P=0.01). No association was found of *B. theta-* and *B. theta+* groups with primary tumor sites (Pearson’s chi-squared test P-values 0.23 and 0.37 respectively).

Supervised hierarchical clustering was used to group the topmost common species according to similar abundance profile across the ordered tumors (Fig. 2E). Similar profiles of a set of species across different conditions would suggest their existence as a community. The clustering delineated 2 major types of the bacterial communities referred as Oral type (O-type) and Gut type (G-type). O-type communities dominated by species that are typical for oral cavity including *Granulicatella adiacens, Rothia mucilaginosa*, and *Leptotrichia unclassified*. G-type community is dominated by known common gut bacteria including *B. theta* and *Akkermansia muciniphila*, well-known known mucus layer degraders in the gut, putative species of *Blautia, Lactobacillus gasserai*, as well as some other bacteria that are not typical for human as the host.

Because the abundance of O-type community varied in *B. theta-* tumors and was decreased in tumors with low richness (Fig. 2F), we further subdivided the tumors into to 2 subgroups (Fig. 2G): Oral High (OH) group with high abundance of oral species and high species richness in the intratumoral microbiome, and Oral Low (OL) group with low abundance of oral species and low species richness. In *B. theta*+ tumors, the abundance of G-type community also varied, but gut species had increased abundance when species richness was low (Fig. 2F). The tumors, therefore, were also subdivided into 2 subgroups: Gut High (GH) group with high abundance of gut species and low species richness in their intratumoral microbiomes, and Gut Low (GL) group with low abundance of gut species and high species richness. The identified 4 subtypes of ACC intratumoral microbiomes, *B*.*theta-* OH and OL and *B*.*theta+* GH and GL, showed significant association (nonparametric MANOVA P= 1e-04 by, and PERMANOVA P= 0.001) with taxonomic composition of the bacterial community (Fig. 2H, Fig. 2I). Species of *Firmicutes, Actinobacteriota*, and *Fusobacteriota*, were more abundant in *B. theta-* negative group (t-test P=0.06, P=0.03, P=0.10 respectively), while species of *Proteobacteriota* was significantly enriched in *B. theta+* tumors (t-test P=0,02), especially in samples characterized by low species richness (GH group) when compared with OH group (P=0.002). Interestingly, species of *Patescibacteria* and *Cyanobacteria*, that are not common inhabitants of the oral cavity, had increased abundances in *B. theta+* tumors (t-test P=0,005 and P=0.06 respectively).

Significant positive correlation of species richness between paired tumor and normal tissue in the study (Fig S3) suggests that dominance of certain species, oral or gut-associated, within the tumor might happen because bacteria of normal tissue colonize the developing tumor of the patient. We, therefore, explore correlations of species abundances between tumor and normal (Fig. 2E) in 33 paired samples to validate the hypothesis. Heatmaps of tumors and normal pairs with the same order of samples and OTUs as in Fig. 2E are presented in Fig. 3A and Fig.3B respectively. Correlation analysis for each species across 33 samples is provided in Fig. 3C. The results support colonization of tumor by oral species from normal tissue, because their abundances in tumors significantly positively correlate with their abundances in normal tissue (mean R=0.65, P<0.02). Abundances of species comprised the gut-type community in tumors were not correlated with normal tissues (mean R=0.02; P>0.1) suggesting *de novo* colonization of the tumor by non-oral, mainly gut-associated species.

**Fig. 3.**
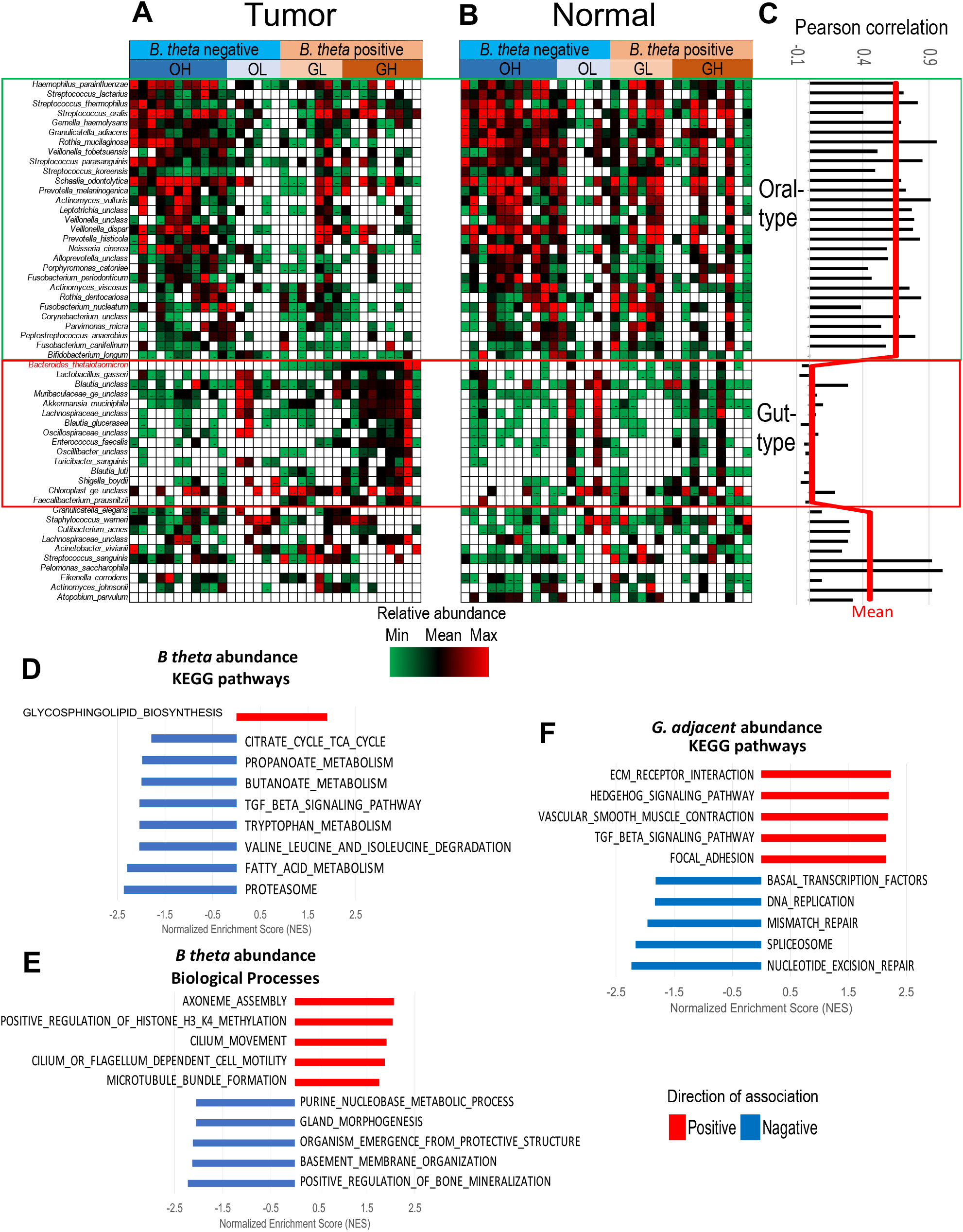
Distinct characteristics of species comprised gut- and oral-type communities in ACC tumors. **A, B**. Heatmaps of most common species in 33 pairs ACC (A) and adjacent normal samples (B). Species and samples in both heatmaps have the same order as in Fig. 2C. Species of the gut-type community are abundant in ACC tumors, but low abundant or absent in their adjacent normal counterparts. **C**. Pairwise Person correlation analysis show significant association of the species abundances between tumor and adjacent normal in the oral-type community and no association for species of the gut-type community. **D, E**. KEGG pathways and Biological Processes in ACC cells significantly associated with the abundance of *B. theta*. **F**.KEGG pathways significantly associated with the abundance of *G. adiacens* in ACC tumors.

### Biological processes correlated with abundances of *B. theta* and *G. adiacens* in ACC tumors

To identify biological processes correlated with abundances of *B. theta* and *G. adiacen*s in ACC tumors, we interrogated our previous RNA sequencing [4] of 44 ACC tumors using GSEA. The analysis (Fig. 3D) identified only one KEGG pathway, biosynthesis of glycosphingolipids, that was significantly positively correlated with *B. theta* abundances in ACC tumors (FDR=0.03).

Glycosphingolipids (GSLs) are known components of the cell membrane. They are comprised of ceramide backbones and glycans that remain outside of the membrane and participate in cell adhesion, cell-cell interaction, and migration. [52-55]. Increased production of GSLs in tumors with abundant *B. theta* suggests that availability of glycans for nutrition may underlie outgrowth of the bacterium. We also highlight activities of related processes including assembly of axoneme, cilium movement, and cilium dependent cell motility (Fig. 3E) [56] among top biological processes positively correlated with B theta. In contrast, activities associated with the normal structure and organization of salivary gland, such as gland morphogenesis and basement membrane organization, negatively correlated with *B. theta* abundances (Fig. 3E).

Several molecular pathways known as significantly more active in ACC-II molecular subtype [4] including extracellular matrix receptors interactions, focal adhesion, Hedgehog, and TGF-beta signaling positively correlated with *G. adiacens* abundance (Fig. 3F). Likewise, processes associated with cell proliferation, a known signature of ACC-I molecular subtype, had negative correlation with *G. adiacens* abundance.

### Comparison of intratumoral microbiomes in ACC with oral and head and neck cancers

The bacterial communities identified in ACC may resemble those found in oral squamous cell carcinoma (OSCC) and head and neck squamous cell carcinoma (HNSC), as all three cancers are originated in or near oral cavity. To identify bacterial genera that the cancers may share with ACC, we examined previously published datasets of intratumoral microbiomes of these cancers. We selected the most common genera from each of the three OTU tables and clustered each table using unsupervised hierarchical clustering with the same parameters (Fig. 4A, Fig. 4B, Fig. 4C). The clustering has identified 2 communities of bacterial *Genera* (Community 1 and Community 2) in each cancer. Many genera were shared between ACC and OSCC (36%) and between ACC and HNSC (34%) and clustered in the Community 1, which is also referred as shared. Importantly, 6 *Genera* of healthy oral microbiome, *Neisseria, Leptotrichia, Actinomyces, Streptococcus, Rothia, and Veillonella*, were shared among all three cancers and clustered close to each other in the shared Community 1. Similar to ACC, this bacterial community was enriched in OSCC and HNSC tumors characterized by a more diverse intratumoral microbiome (Figs. 4D and 4F, respectively), by a less aggressive phenotype (Figs. 4B and 4C, respectively), and by better overall patient’ survival (Figs. 4E and 4G, respectively). Thus, the shared oral genera may be considered as biomarkers of healthier tumor microbiome in all three cancers. In contrast, Community 2 was comprised of distinct bacteria in each of three cancers and associated with more aggressive phenotype and worth patient’ survival.

**Fig. 4.**
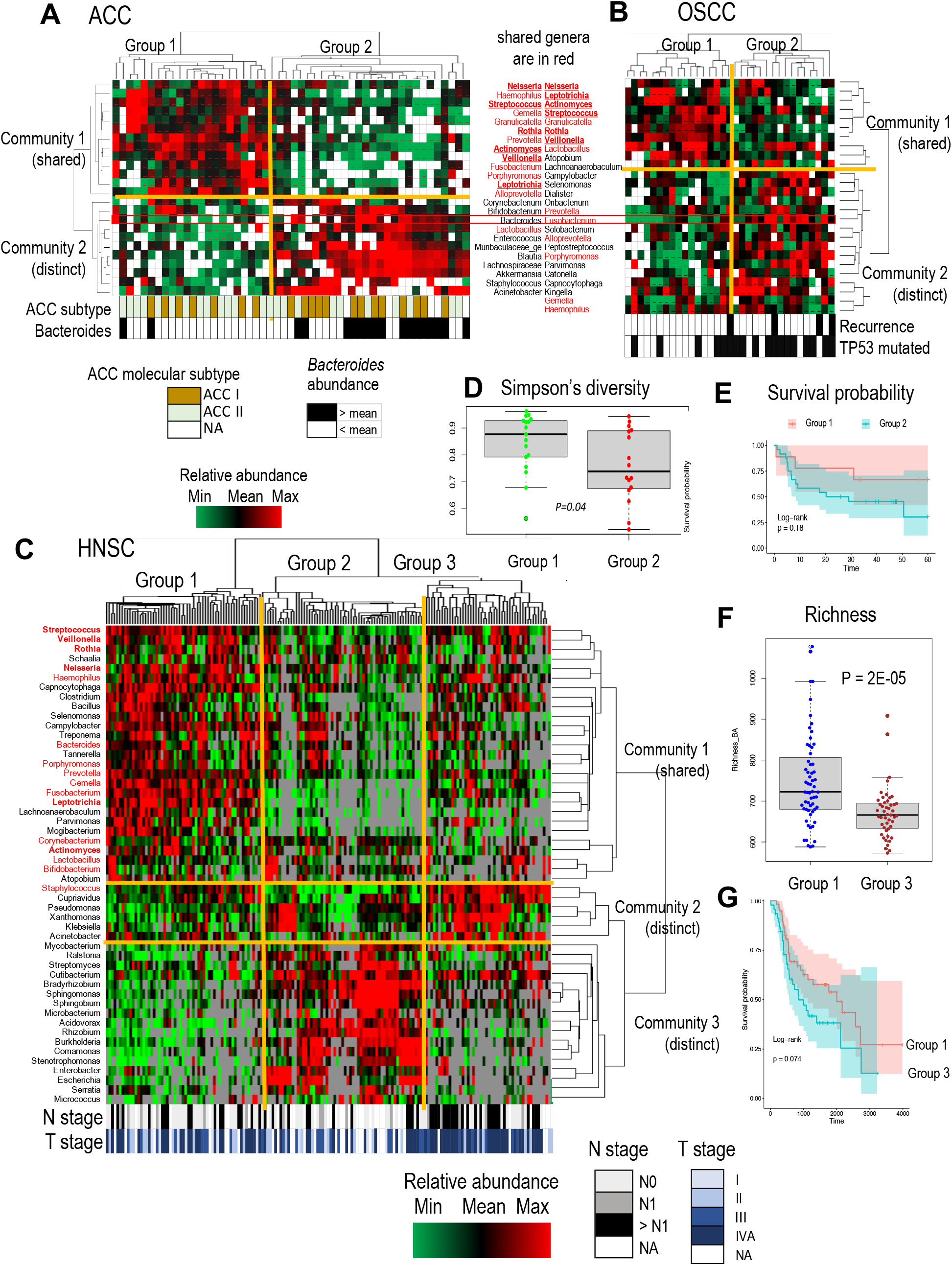
Comparative analysis of intratumoral microbiomes in ACC with OSCC and HNSC. A, B, C. Unsupervised hierarchical clustering of tumors in ACC (A), OSCC (B), and HNSC (C) identified 2 communities of bacteria (rows) in each cohort: Community 1 populated by shared oral *Genera* (names are colored in red) and Community 2 populated by *Genera* that are mainly specific for each cohort (names are colored in black). Names of Genera identified in all three cohorts are given in red bold color. Columns of each heatmap denotes tumors that are clustered together into the major tumor groups (top of each heatmap) with similar abundance profile of the intratumoral communities. Bars at the bottom of each heatmap show characteristics associated with the tumor groups. In ACC (A), the clustering identifies 2 tumor Groups that recapitulate the supervised clustering (Fig. 2B and 2E) based on intratumoral *B. theta* abundance. Group 1 ACC tumors have many healthy oral Genera, but less abundant *Bacteroides* (t-test P=0.0006) and other gut-associated *Genera*. The aggressive ACC I molecular subtype is also less common in patients of the group (Fisher’s test P=0.14). In OSCC (B), Group 1 tumors have less abundant *Fusobacterium* (t-test P=0.0001), significantly reduced frequency of TP53 mutations (Fisher’s test P=0.037) and of disease recurrence (Fisher’s test P=0.044). *Bacteroides* and *Fusobacterium* are labelled by red rectangles in ACC and in OSCC respectively as potential biomarker of more aggressive phenotype. In HNSC (C), the clustering identifies 3 tumor groups (Group 1, Group 2. and Group 3). Group 1 HNSC tumors is less likely than Group 3 have advanced stage (Fisher’s test P=0.01) and a spread of the cancer into lymph nodes (Fisher’s test P=0.05). No difference was found between Group 1 and Group 2 tumors. **D, E**. Community 1 in OSCC was associated with more diverse microbiome (D) and better overall patients’ survival (E). **F, G**. Community 1 in HNSC was also associated with more diverse microbiome (F) and better overall patients’ survival (G).

## DISCUSSION

The study reveals two major microbial subtypes, oral-like and gut-like, in an enclosed salivary ACC (Fig. 5A). They are characterized by different microbial communities (oral or gut type), different tumor phenotype (less or more aggressive), association with different molecular subtype of ACC (ACC-II or ACC-I), and by better or worth probability of overall survival. Patients with increased number of bacterial species (high richness) in normal tissue have oral type normal and tumor microbiomes that are similar in taxonomic composition and populated by many shared oral taxa, such as *Granulicatella, Rothia*, and *Leptotrichia*, positively associated with the patient survival. Because submandibular and minor salivary glands are main sites of ACC in the cohort (62% of samples), it is expected that bacterial communities of the glands are shared with the oral cavity. In a published study of bacterial microbiomes from healthy humans [57], oral sites had high relative abundance of *Streptococcus, Veillonella, Prevotella, Neisseria, Actinomyces, Leptotrichia, Rothia*, and *Porphyromonas*. They are the same genera found to be abundant in the oral type of microbial community in ACC tumor and normal tissues (Fig. 4B, 4C, 4D). Furthermore, a comparative analysis of ACC intratumoral microbiomes implemented in this study with OSCC and HNSC linked 6 of these oral genera to more diverse intratumoral community, better survival, and to less aggressive tumor phenotype in all three cancers. The findings suggest co-occurrence and interaction of the oral genera leaving as a bacterial community and highlight their potential role in tumor suppression not only in ACC, but also in other cancers associated with the oral cavity. The positive effect of the oral community on survival can be due a favorable tumor microenvironment supporting diversity, functional redundancy, resilience and stability of not only the microbial genera but also host immune cells in the microenvironment [46, 58-60]. In addition to these tumor suppressive effects, there are likely more specific molecular mechanisms. In our study, the most significant positive effect on survival in ACC was found for *Granulicatella adiacens*, a common inhabitant of normal oral microbiota, The effect can be explained by nutritional requirements of this fastidious bacterium that requires vitamin B_6_ (pyridoxine) for growth [61]. Because pyridoxine must be supplied from the environment, increased intake of the vitamin by growing *G. adiacens* may reduce availability of *B*_*6*_ for proliferating ACC cells [62]. Further experimental research is needed to provide evidence of *G. adiacens* involvement in ACC pathogenesis and to explore specific molecular mechanisms.

**Fig. 5.**
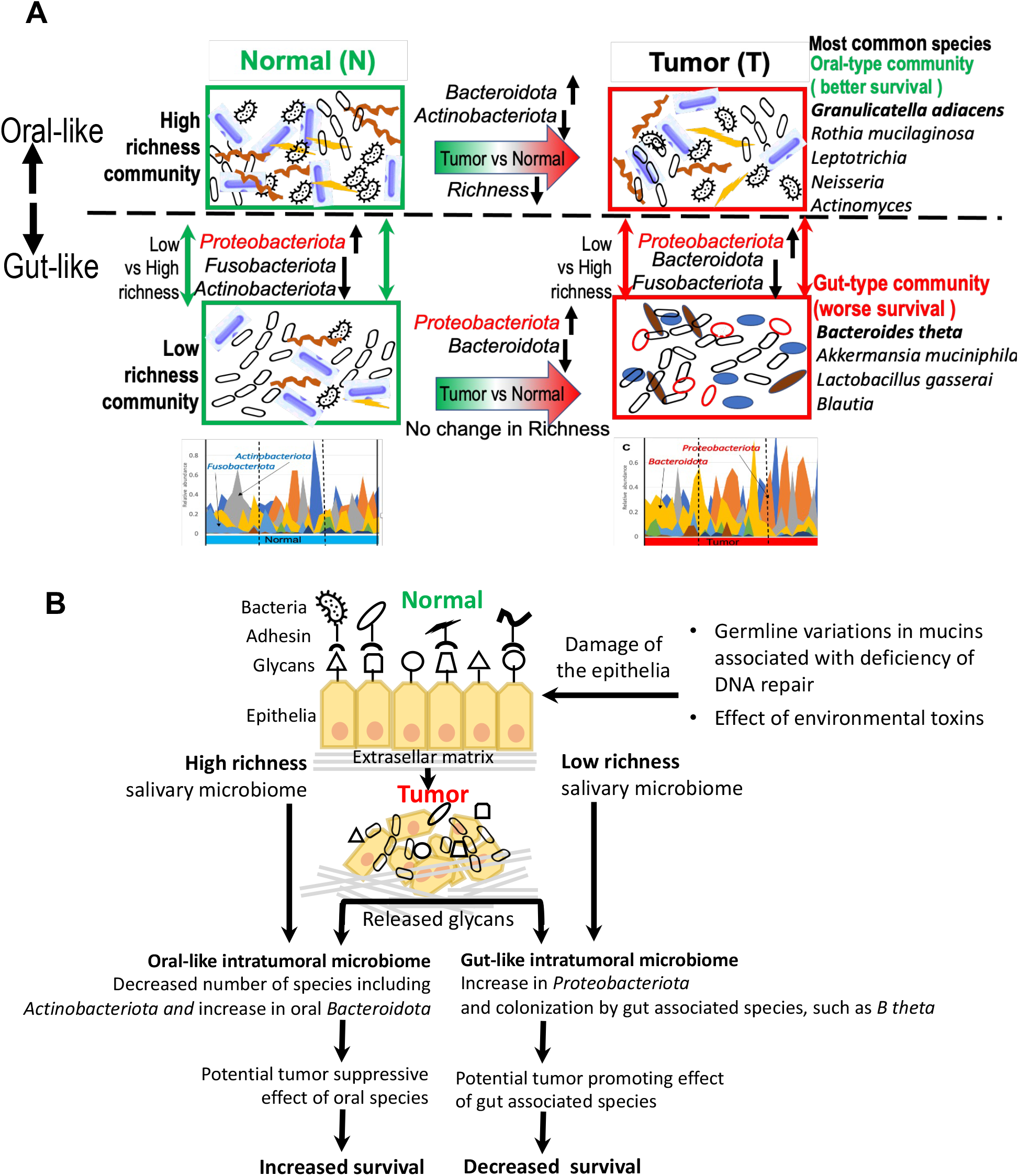
Graphical summary of the results. **A**.Distinct taxonomic structure of intratumoral microbiome in ACC with high and low species richness and association with survival. Rich microbial community in normal tissue is linked to oral type intratumoral community dominated by oral species with significantly increased abundances of *Bacteroidota* and decreased abundances of *Actinobacteroidota*. Microbial community with low richness is linked to gut type intratumoral community populated by many species involved in mucus layer degradation. **B**.Hypothetical model linking ACC development and progression to distinct changes in taxonomic composition of intratumoral microbiome and feedback effects. Sustain damage of salivary epithelia in course of ACC development leads to releasing of salivary glycans and outgrowth of glycan degrading bacterial taxa. In low richness intratumoral microbiome, which is not resilient, the outgrowing taxa are represented by non-oral gut-associated bacteria and *Proteobacteriota* promoting tumor progression. In high richness intratumoral microbiome, which is more resilient, the taxa are represented by oral *Bacteroidota*. Althoug most other oral species are decreased, the colonization by gut associated glycan degraders is suppressed by the oral microbes.

The gut-like microbial subtype was found in ACC tumors with reduced species richness in normal salivary tissue. The subtype is associated with dramatic changes in taxonomic structure of intratumoral microbiomes suggesting an outgrowth of taxa that are not typical for the oral cavity. Multiple new colonizes of the communities are common inhabitants of the human gut, including *B. thetaiotaomicron* that have significant negative association with the patient survival. Although low species richness was not linked to a distinct taxonomic structure of the intratumoral microbiome in any cancer, abundant gut species and *Proteobacteriota* observed in ACC tumors with low richness are not surprising and can be explained by low resistance of such microbial communities to colonization by other species including pathogens [58, 63]. Indeed, *Proteobacteria*, was significantly expanded in low richness ACCs. The *Phylum* is a known microbial signature of human diseases and impaired epithelial tissue [64-66]. It also has strong association with metastasis and poor survival in cancer patients [67]. Other uncommon ACC colonizers including gut microbes from genera of *Bacteroides, Akkermansia, Blautia, Enterococcus, Faecalibacterium, Fusobacterium, Bifidobacterium* [57, 68] also can be linked to metabolic and structural changes in tumor tissues [69]. Several studies reported not only presence, but also involvement of oral microbes in pathogenesis of gastrointestinal inflammatory diseases [70], [71], and of colon cancer [72]. It has been demonstrated that a disease associated metabolic shifts in the gut may drive the colonization of gut by oral *Veillonella parvula* or *Fusobacterium nucleatum*. In our study, we observe a reverse oral-gut microbiome interaction, a colonization of low richness ACC tumors by gut microbes. There is also a possibility that other occasional non-human inhabitants, such as plant associated *Bradyrhizobium centrosematis* [73] can colonize the low richness tumors. The bacterium is also a common pollutant of the drinking water [49] and has been reported as NGS contaminants [74, 75]. Our study, however, found *B. centrosematis* as differentially abundant only in tumors with low number of species but not in normal tissue or in tumors with rich microbiome. Therefore, expansion of the bacterium in the tumors may be relevant to ACC pathogenesis. Additional experimental studies are needed to confirm that.

Notably, several intratumoral bacteria, including *Akkermansia, Bacteroides, Blautia, Bifidobacterium*, and *Enterococcus*, found in low richness ACCs are not only known human gut colonizers, but also can utilize mucus glycans for their growth [76]. In high richness ACCs, oral species of *Bacteroidota*, known by consumption of mucus glycans, also significantly increased their abundances in comparison with matched normal tissue. We propose, therefore, that a release of glycans by developing tumors may lead to expansion of the glycan degraders (Fig.5B). Glycan liberation in ACC can result from sustained damage to the salivary epithelia due to DNA repair pathway deficiencies and increased frequency of rare germline mutations in mucins that defend epithelial tissue against toxins [77]. Consistent with the hypothesis, the GSEA implemented in the study for tumor RNA has identified biosynthesis of the cell membrane components comprised of glycans as significantly positively correlated with *B. theta* abundances and activities associated with normal organization of salivary gland as negatively correlated (Fig. 3D, 3E). Expansion of glycan-degrading bacteria may promote the development and progression of ACC due to positive feedback effects on tumor growth. Degradation of salivary glycans damages the mucus layer that protects epithelial cells from the environment and helps to maintain the healthy tissue of the salivary gland [22, 78]. The degradation may also produce byproducts, such as ornithine and ceramides, with tumor promoting effects [79]. In addition, monosaccharides released as byproducts of the mucus degradation [80] can lead to growth and reprogramming of tumor cells. Further studies are necessary to prove the tumor promoting effects of *B. theta* and other mucus degraders and reveal underlying molecular mechanisms.

## CONCLUSIONS

Two microbial subtypes dominated by oral versus gut bacterial species were identified in ACC. The oral-like subtype is associated with less aggressive tumor phenotype, ACC-II molecular subtype, and better probability of the patient survival. *Genera* dominated in the subtype are shared with the oral community found in healthy individuals and with OSCC and HNSC intratumoral microbiomes. In both cancers, intratumoral communalities with the shared oral species were associated with more diverse microbiome, less aggressive tumor characteristics, and better survival. The gut-like ACC microbial subtype is characterized by increased abundance of known mucus layer degraders dominated by gut species in low richness microbiomes and by oral *Bacteroidota* in case of high richness. Most significant negative association with the survival in ACC was observed for *B. theta*, likely because of damage and reprogramming of salivary epithelia in course ACC progression resulting in release of glycans consumed by the organism.

## Acknowledgement

We would like to acknowledge the High-Performance Computing for research facility (HPCC) and Platform for Innovative Microbiome & Translational Research (PRIME-TR) at the University of Texas MD Anderson Cancer Center for providing resources contributed to the research results reported in this paper.

## Availability of Data and Materials

The 16S rRNA gene sequencing data were deposited into the NCBI Sequence Read Archive (http://www.ncbi.nlm.nih.gov/sra) under the BioProject accession number PRJNA1049986.

## Author contributions

Study design: TK, AE, YM

Samples preparation and analysis: YM, CC, IF, LM, YH, FV

Data processing: TK, XW, XS, CC

Supervision: JZ, AE, NA, RJ, AF Validation: TK, NA, RJ

Writing - original draft: TK, AE

Writing - review and editing: RF, YM, LC, NA, RJ, JZ, AF

## Supplementary Information

Additional file 1. Supplementary Figures 1-10.

Additional file 2. Supplementary Tables

Table S1. Clinical characteristics of the cohort

Table S2. Cox Proportional-Hazards Model (coxph) output for association of overall survival with clinical characteristics, measures of diversity, and topmost abundant taxa.

Table S3. Cox proportional-hazards model identifies B. theta abundance as an independent predictor of survival from clinical characteristics and measures of diversity.

