## Additional file 1. Supplementary Figures 1-10 for "Intratumoral Microbiome of Adenoid Cystic Carcinomas and Comparison with other Head and Neck Cancers"

Fig. S1 . Study overview. Red border indicates data set generated in this study.

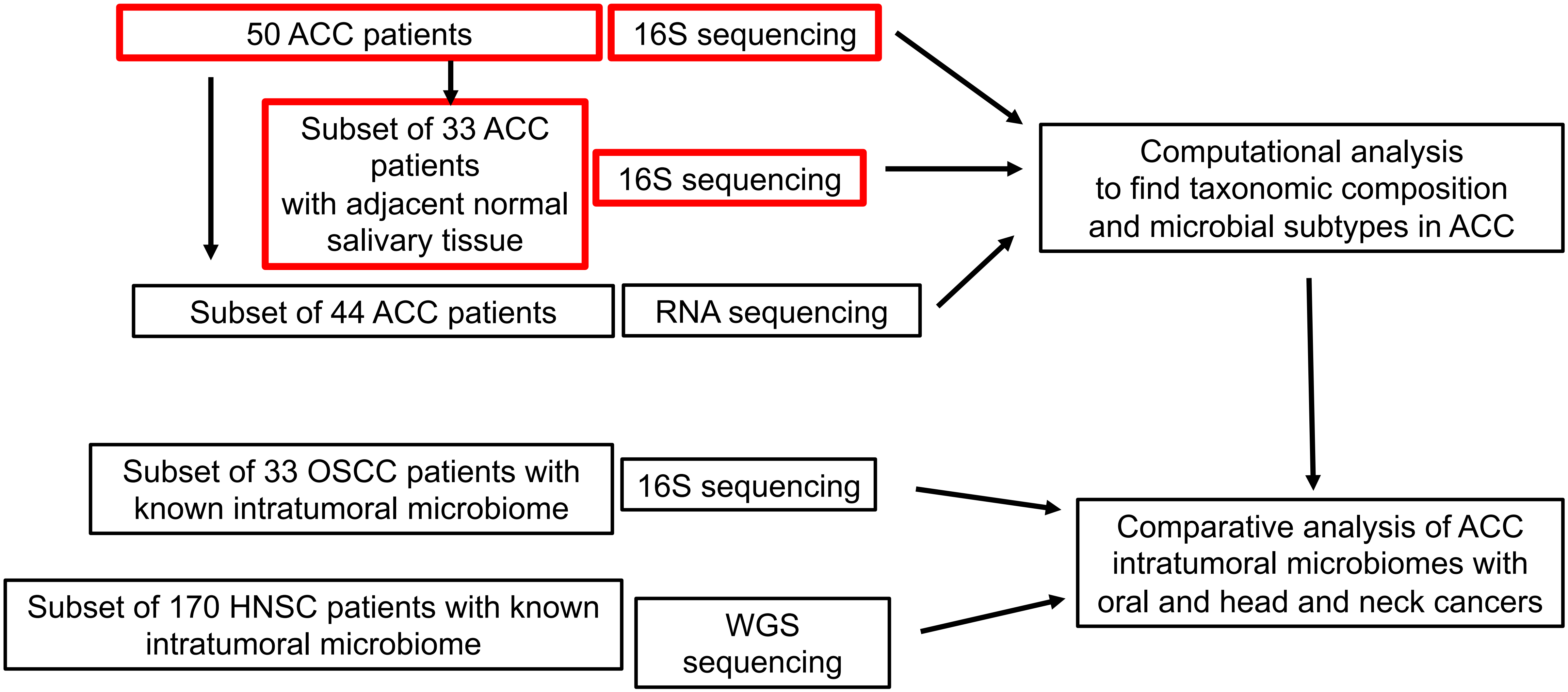

Fig. S2. Evaluation of the batch effect. Unsupervised hierarchical clustering of samples in terms of most common OTUs reveal similar abundance profiles for 5 duplicated/triplicated samples. The samples are clustered near each other. There is no significant difference between Cluster 1 and Cluster 2 in terms of the batch assignment (Fisher's test p-value p=0.12)

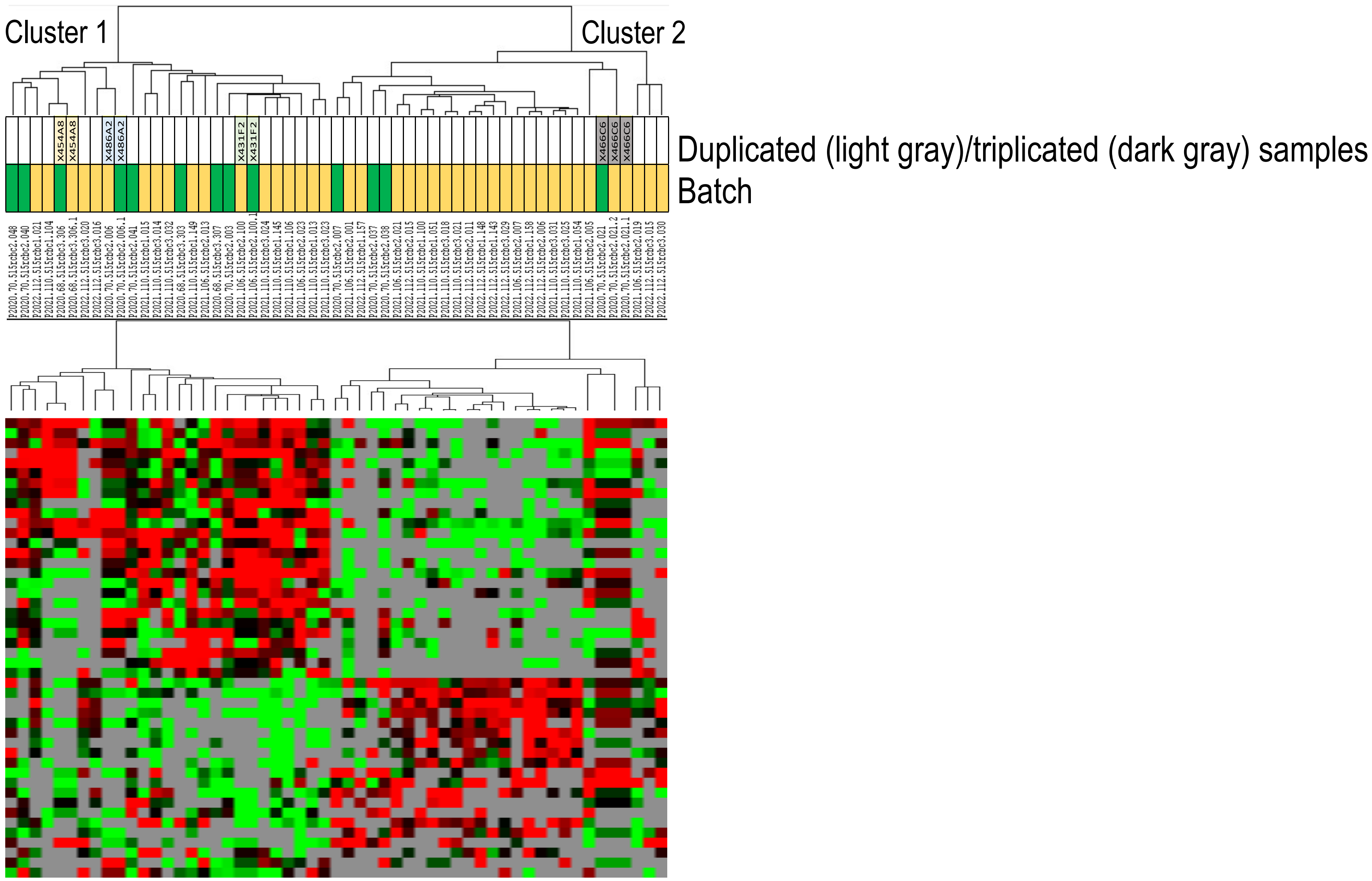

Fig. S3. Significant correlation between richness in normal and tumor tissue.
Red dashed line indicate perfect relationship. Blue dots in the scatter plot mark samples with a specific richness values in tumor and normal.

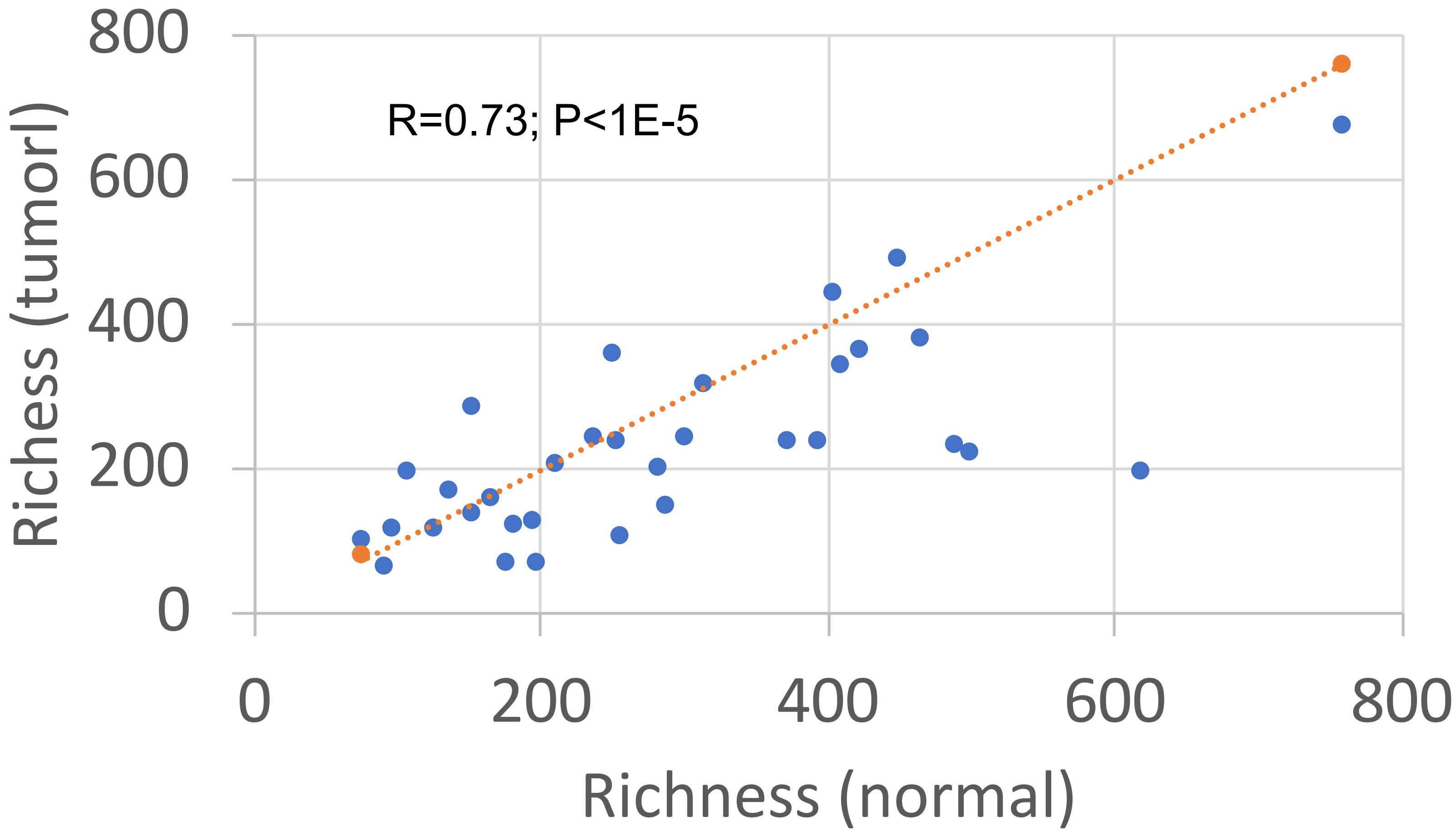

Fig. S4. Permutational multivariate analysis of variance of 33 paired tumor-normal tissue in terms of bacterial phyla.
There is no difference between tumor and normal samples if richness of  samples are not considered (A) because the characteristic significantly effect taxonomic structure of bacterial communities in terms of phyla  (B). Difference between tumor and normal samples becomes more significant If only samples with rich microbiomes are considered (C), but not for samples with low low richness microbiomes (D).

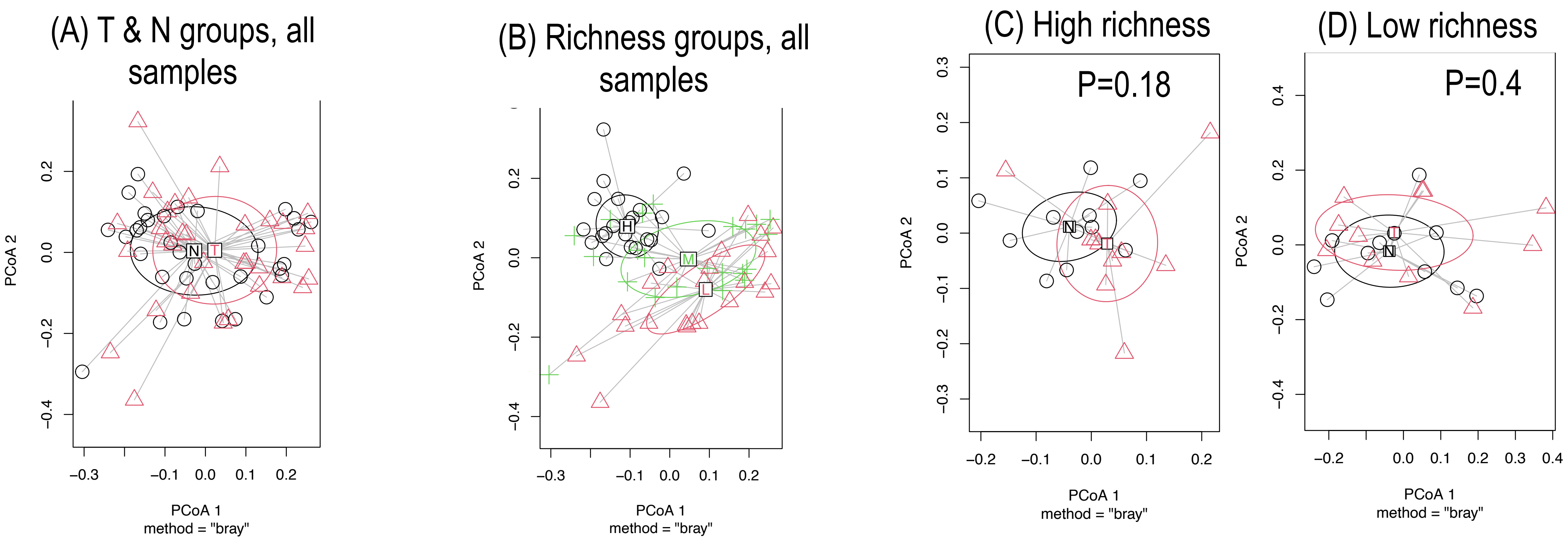

Fig. S5. Decreased abundance of *Actinobacteriota* in normal tissue and *Bacteroidota* in tumors with low richness.

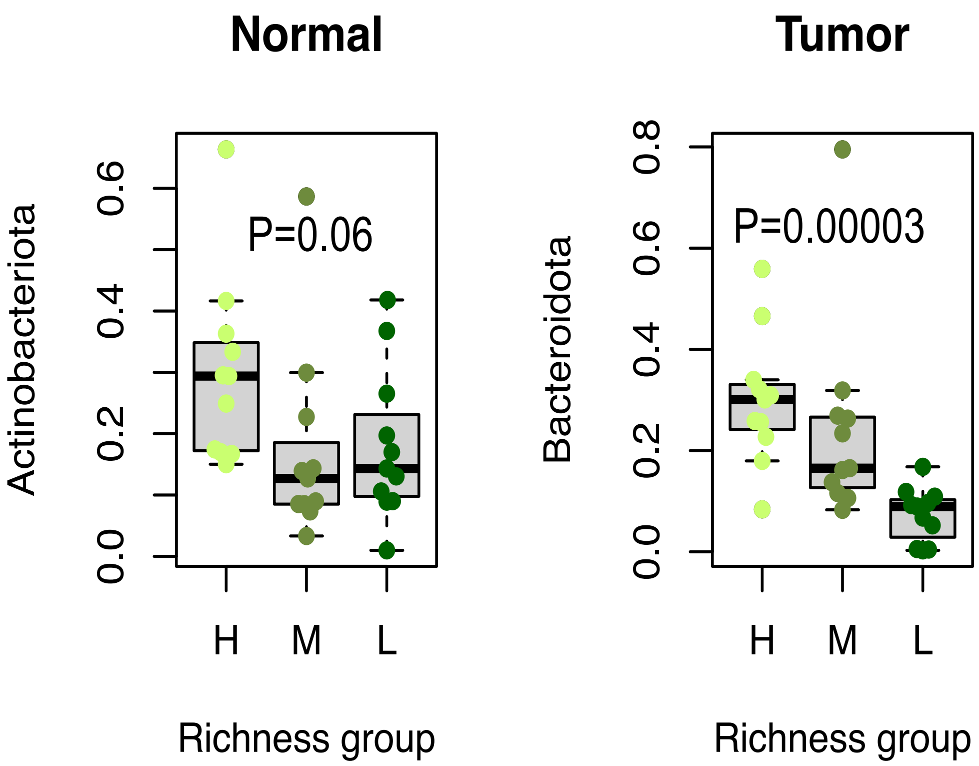

Fig. S6. Increased abundance of *Proteobacteriota* in tumor vs normal tissue in low richness group of samples.

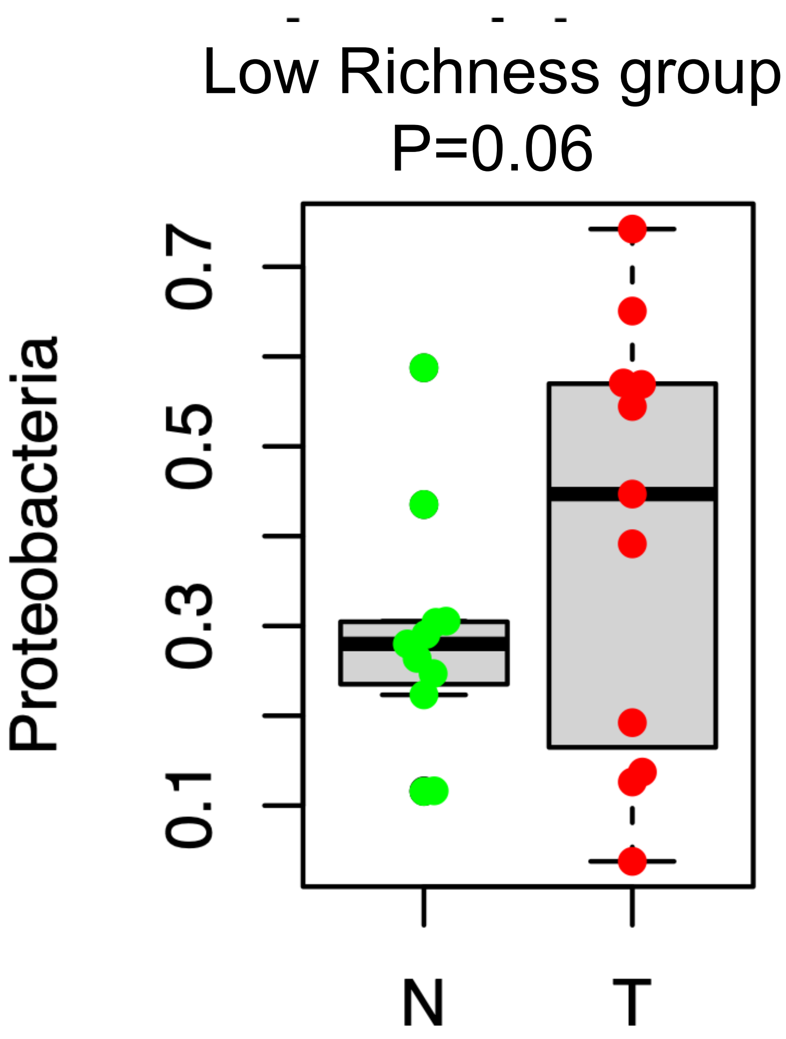

Fig. S7. Relative abundance of top *Orders* in 33 paired tumor-adjacent normala samples.

Abundances of spp at the order level were sorted by p-value (from low to high) and only those that have mean relative abundance more than 0.009 and p<=0.1 were selected. Normal samples are sorted by Richness (from high to low), and tumor samples have the same order as normal. Order *Xanthomonadales* (class *Gammaproteobacteria*) and orders of bacterial pathogens, *Oceanospirillales* and *Rhodobacterales*, were more abundant in tumor tissue (P=0.03, P=0.07, P=0.06 accordingly), while orders of oral bacteria, such as *Pasteurellales* (p=0.02), *Fusobacteriales* (p=0.07), *Lachnospirales* (p=0.09), and *Actinomycetales* (P=0.02, P=0.07, P=0.09,P=0.10 accordingly) were more abundant in normal tissue, especially in samples with high species richness.

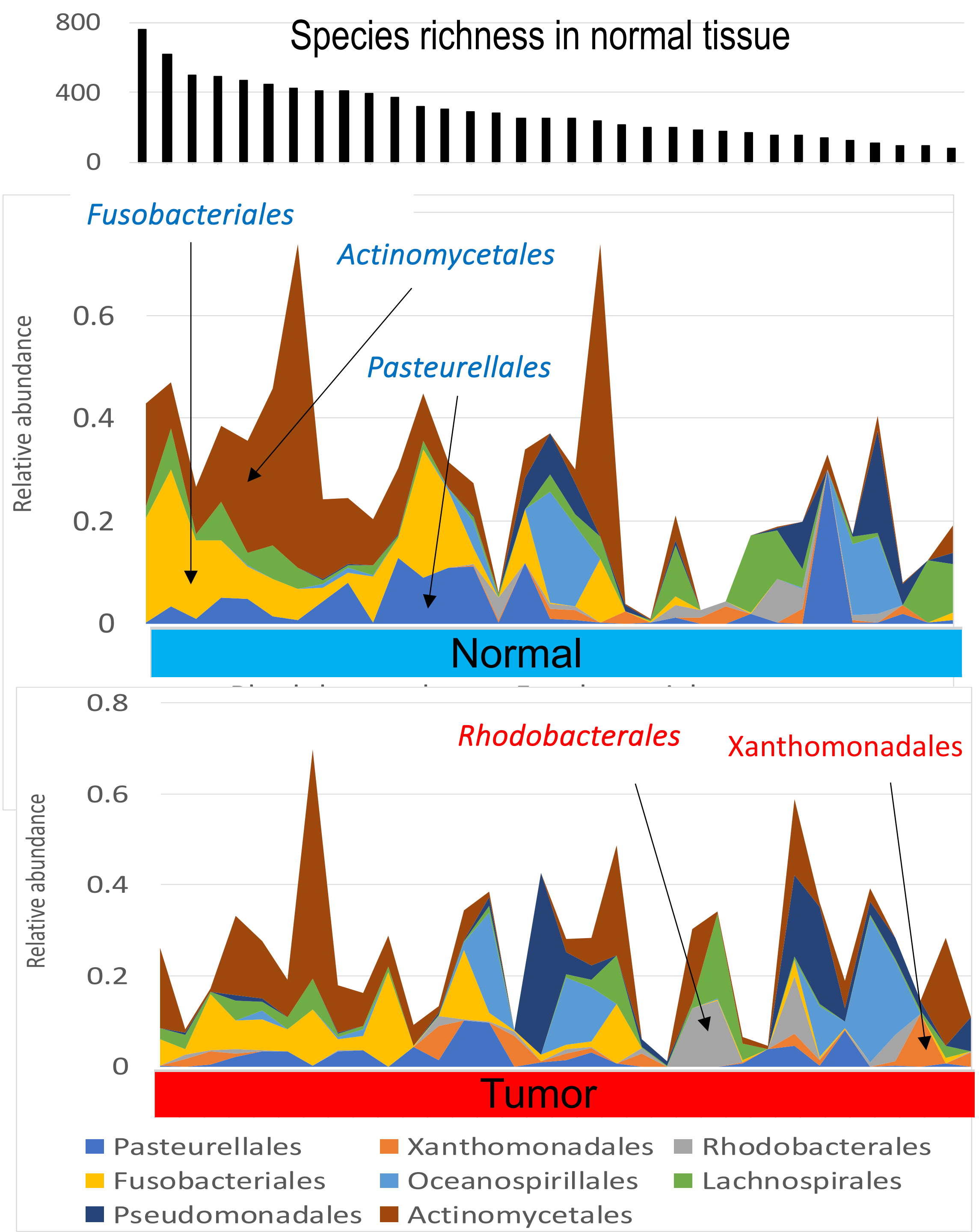

Fig. S8. Differentially abundant taxa between tumor and adjacent normal tissues revealed by LDA Effect Size (LEfSe) analysis. Default parameters was used for the analysis except LDA, which was set to 1.5 to reveal more differentially abundant taxa. Species that were also identified by MaAsLin2 (Fig. 1D) are in red and green rectangles.

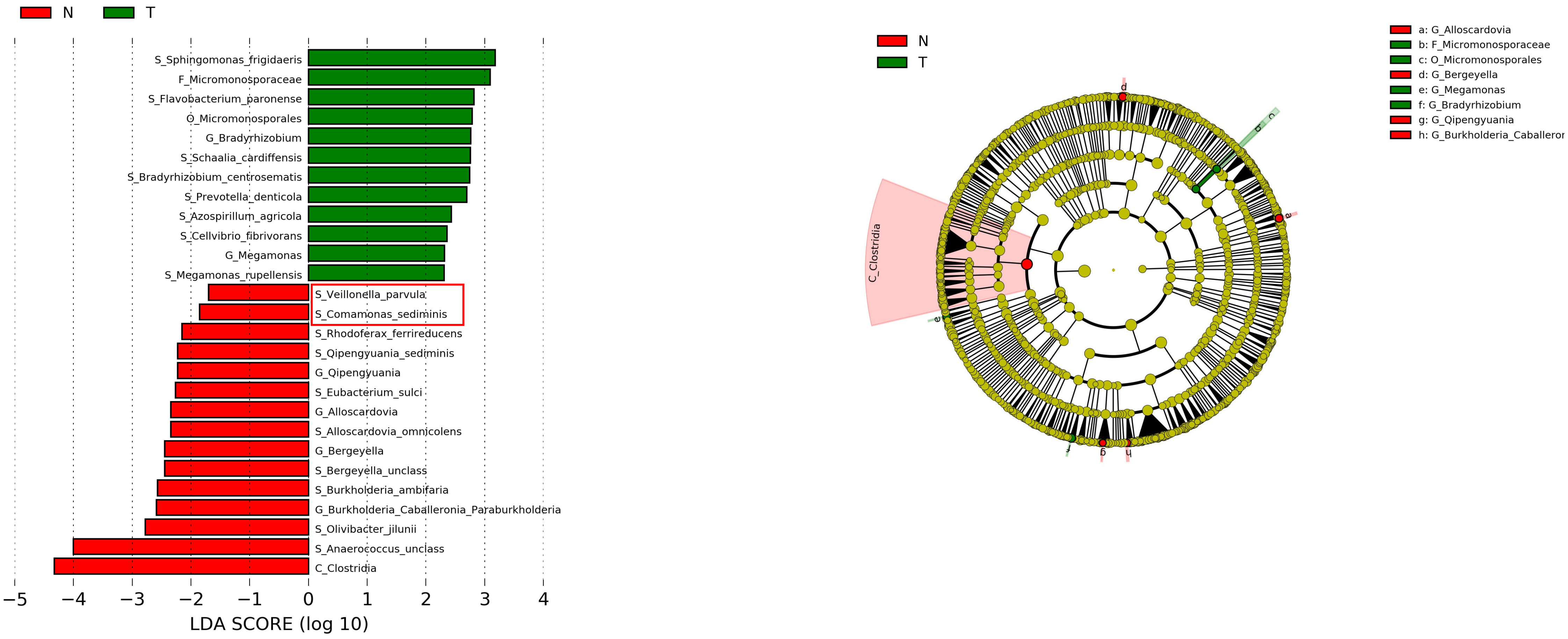

Fig. S9. Abundances of species differentially enriched in normal tissues. Samples are sorted by richness of their microbiomes. The figure shows that species species differentially enriched in normal tissues are mainly found in microbiomes with high richness.

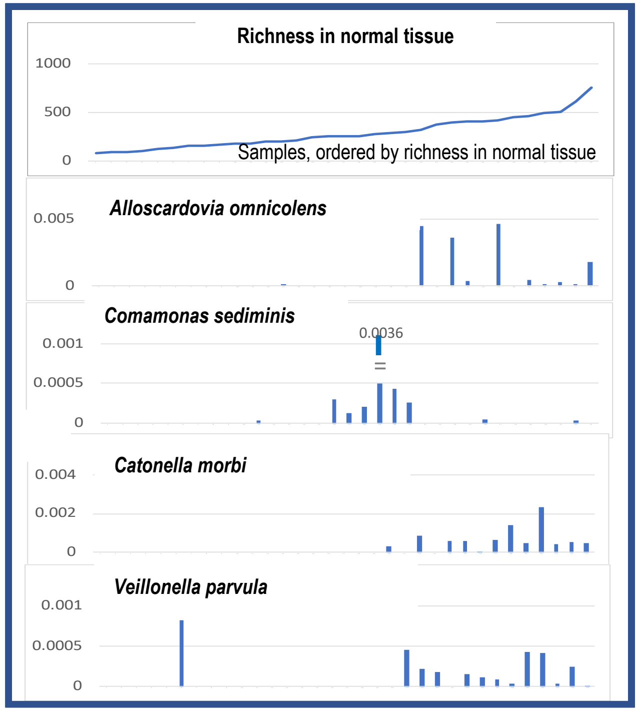


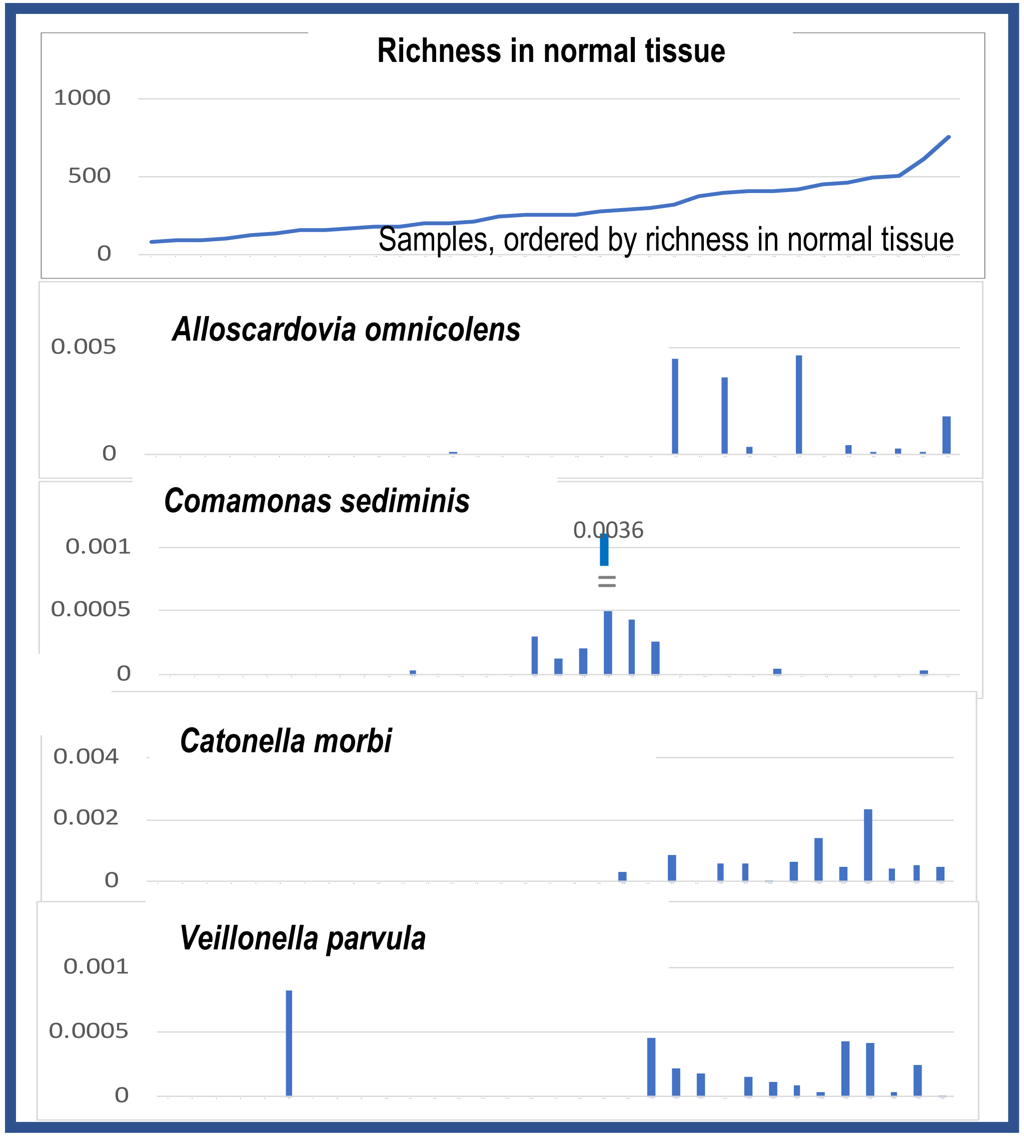

Fig.  S10. Abundances of species differentially enriched in ACC tumors. Samples are sorted by richness of their microbiomes. *Megamonas* and *Bradyrhizobium centrosematis* are mainly found in microbiomes with low richness.

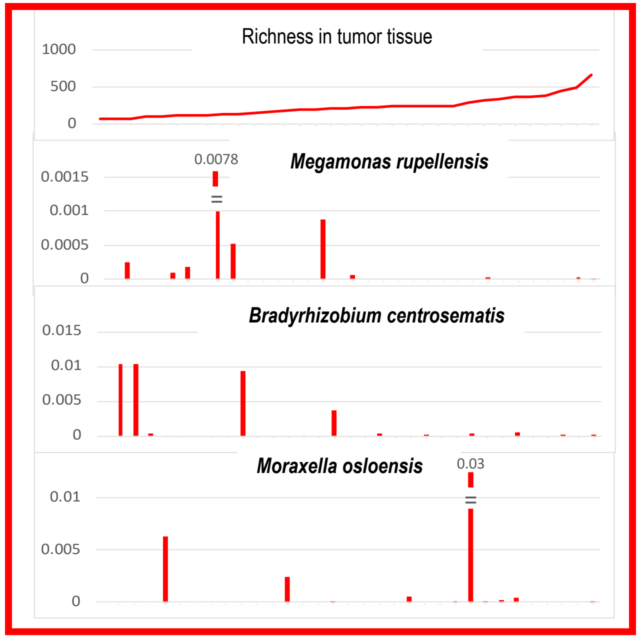


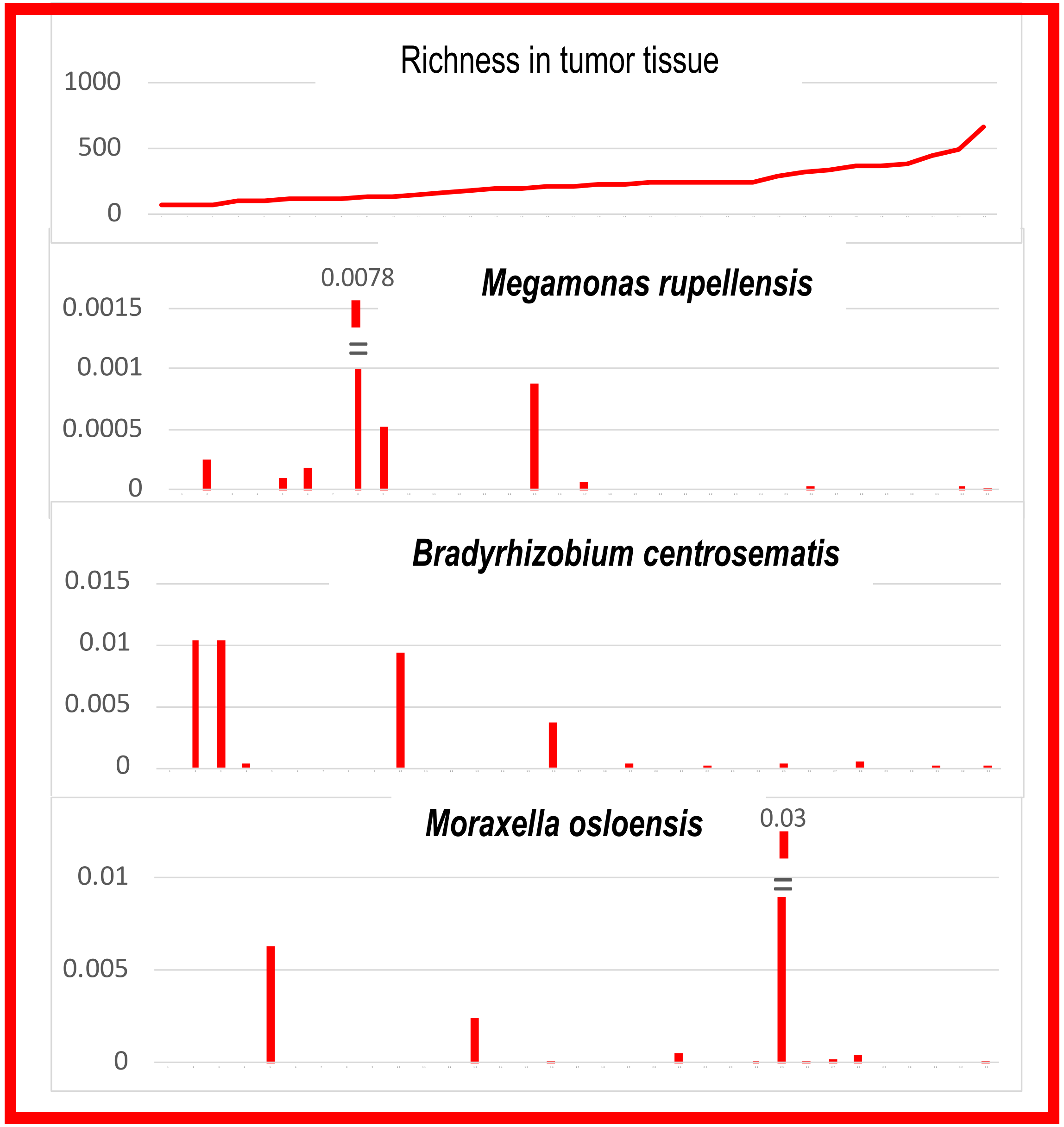
